# The discovery of mixed colonies in *Temnothorax* ants supports the territoriality hypothesis of dulotic social parasite evolution in myrmicine ants

**DOI:** 10.1101/2023.08.08.552493

**Authors:** Sarah Bengston, Anna Dornhaus, Christian Rabeling

## Abstract

Social parasitism, where one social species parasitically depends on the other for survival and reproduction, is a highly successful life history strategy, especially in the eusocial Hymenoptera. In ants alone, more 400 species of socially parasitic species exist and multiple forms of social parasitism evolved independently and convergently. Yet disentangling the evolutionary history of obligate social parasitism is challenging. Identifying species that inform the transition from eusocial toward socially parasitic behavior is crucial for understanding the underlying co-evolutionary processes. Here, we report the first case of mixed colonies involving four predominantly free-living *Temnothorax* ant species from the western United States. Three *Temnothorax* species supplement their worker force with brood from the nests of their four congeners. We suggest, based on these observations and other published evidence, that this facultative dulotic behavior may have resulted from territorial contests due to limited nest sites. Socially parasitic behavior is not present in all populations across the species distribution ranges, however in populations where this behavior was observed, it is also associated with significant increases in interspecific aggression. These four species of Western US *Temnothorax* ants represent a particularly interesting case of social parasitism, because the presence of between-population behavioral variation provides a powerful system to test hypotheses about the ecological and behavioral conditions underlying the evolutionary transition from eusocial to socially parasitic behavior.

## Introduction

Interactions between species are strong drivers of evolutionary change. The co- evolutionary forces between predators and prey, parasites and hosts, and mutualists, for example, are known to shape the evolution of complex traits. In addition, co-evolutionary interactions are known to result in speciation and diversification of organisms in some instances (Thompson 2005). In the case of predator-prey or host-parasite associations, long-lasting and highly specialized adaptations frequently evolve as a result of the escalation of an evolutionary arms race and the resulting reciprocal selection (Dawkins and Krebs 1979; Davies et al. 1989; Hanifin et al. 2008; Kleeberg et al. 2015).

Additionally, the extent of these adaptations may vary across populations as a result of biotic and abiotic environmental factors (Nash et al. 2008), such as population density or abundance (Boersma et al. 1998; Reznick et al. 2001; Cousyn et al. 2001; Bell 2005). For example, if a defense against a parasite is costly, it should only evolve in populations under strong parasitic selection pressure, or high parasite density (Sheldon and Verhulst 1996; Wade 2007; Foitzik et al. 2009; Feeney et al. 2012). Therefore, while social parasitism can only be identified in a single population, it is informative to compare parasitized with unparasitized host populations to better understand the dynamics under which adaptations against parasitism evolve in the host.

Host-parasite interactions have most commonly been approached by applying concepts and approaches from the study of disease and immunology (Yazdanbakhsh et al. 2002; Schmid-Hempel 2011). Studying host-parasite coevolution also significantly influences the fields of ecology and evolution and has allowed us to make significant progress in understanding the evolution of host-parasite arms races, i.e., the continual diversification of parasitic strategies and host defenses (Werren et al. 1995; Sheldon and Verhulst 1996; Poulin 2007). At the macroscopic level, understanding how hosts and parasites interact provides an opportunity to understand the evolution of sophisticated behavioral strategies and the resulting population dynamics associated with trait coevolution. The defense against brood parasites in birds is a particularly prominent example (Davies et al. 1989; Langmore et al. 2003; Davies 2010). Host bird species have developed increased brood discrimination abilities to detect parasite eggs in their nest (Feeney et al. 2012), show different patterns of interspecific aggregation preference in areas with parasites (Clark and Robertson 1979), and increased aggressive and mobbing behavior in parasitized populations (Uyehara and Narins 1995; Welbergen and Davies 2009).

One particularly interesting form of parasitism is social parasitism in the eusocial Hymenoptera. Social parasites are known in all groups of eusocial Hymenoptera (wasps, bees, and ants; Buschinger 1986, Choudhary et al. 1994, Schmid-Hempel 1998, Martin et al. 2010, Michener 2007), and among the approximately 14,000 species of ants alone, at least 400 species of social parasites are known (Buschinger 1986b, 2009; Hölldobler and Wilson 1990a; Rabeling 2021). Social parasites, defined as members of a social species that parasitically depend on another social species, usurp social insect colonies to realize their own fitness (Bourke and Franks 1991). In the eusocial Hymenoptera, social parasitism is remarkably diverse and in ants alone, social parasitism evolved at least 91 times independently (Gray and Rabeling 2023). Since Wasmann’s and Wheeler’s (Wasmann 1891, 1909; Wheeler 1904, 1919) earlier studies on socially parasitic ants, the following three categories have generally been adopted to roughly characterize the convergently evolved life-history strategies of socially parasitic ants (see also, for example: Emery 1909; Wheeler 1910; Wilson 1971; Buschinger 1986a, 2009; Hölldobler and Wilson 1990b; Brandt et al. 2005; Huang and Dornhaus 2008; Rabeling 2021). (1) Temporary social parasitism: The parasite queen invades the host colony, kills the host queen, and produces her offspring, which is cared for by the host workers. After colony founding host and parasite workers are found inside these same colony, but eventually host workers die off leaving only the established colony of the parasitic species (Hölldobler and Wilson 1990b). (2) Dulotic social parasitism: Foundress queens of dulotic species establish colonies via temporary social parasitism. In mature colonies, workers of dulotic species ‘raid’ other host colonies for pupae, which are then in part eaten whereas some eclose in the parasite’s colony. These host workers forage, care for the parasite’s brood, and in some cases feed the parasites. The interaction between dulotic species and their hosts may be facultative or obligatory (Mori et al. 2001). Two primary hypotheses have been proposed for how this form of social parasitism may have evolved.

Darwin proposed that predatory ants would raid other nests for brood and some of the stolen brood would subsequently eclose before they were eaten (Darwin 1859; Alloway 1980; Pollock and Rissing 1989; Borowiec et al. 2021). Alternatively, the ‘territoriality hypothesis’ was proposed suggesting that intraspecific territorial contests can induce selection for brood raiding (Stuart and Alloway 1982). This facultative relationship between rival colonies may then facilitate the obligate evolution of dulotic social parasites and their hosts (for a further review, see Pollock and Rissing 1989). (3) Inquiline social parasitism: The parasite queen resides in the nest of the host and is cared for by the host’s workers. In most cases the parasite queen does not produce workers.

Host queen tolerant inquilines co-exist with the host queen inside the same colony, where the host queen continues to produce workers. The parasite queen often inhibits the host queen’s production of sexual offspring (Buschinger 1990; Hölldobler and Wilson 1990b; Bourke and Franks 1991; Rabeling and Bacci 2010). Some species of inquiline social parasites were shown to evolve reproductive isolated from their hosts in sympatry (Savolainen and Vepsäläinen 2003; Rabeling et al. 2014; Mera-Rodríguez et al. 2023), and therefore it is of particular interest to understand the evolutionary origin of inquilinism.

The myrmicine ant genus *Temnothorax* is an interesting study system for exploring the behavioral ecology of socially parasitic ants, because both dulosis and inquiline social parasitism occur frequently in this genus (Alloway 1980; Herbers and Foitzik 2002; Beibl et al. 2005; Prebus 2017). The genus *Temnothorax* is highly speciose with 500 described species and subspecies [Note: The exclusively parasitic satellite genera *Protomognathus, Myrmecoxenus,* and *Chalepoxenus* were synonymized under the name *Temnothorax* (Ward et al. 2015)], which are distributed throughout the world (Bolton 2018). Most *Temnothorax* species are small ants (2-3 mm) that often live in pre- formed cavities in rocks, acorns or twigs (Rüppell et al. 2003; Rueppell and Kirkman 2005; Pamminger et al. 2012; Bengston and Dornhaus 2013). The genus has a worldwide distribution, though of the species in the United States, more is known about the natural history of the eastern species than the western species (Fisher and Cover 2007, Foitzik et al. 2004, Achenbach and Foitzik 2009, Modlmeier and Foitzik 2011, Pamminger et al. 2012; but see Rüppell et al. 2003; Rueppell and Kirkman 2005; Bengston and Dornhaus 2013, 2015; Heinze and Rueppell 2014).

Here, we describe a new instance of facultative social parasitism in western *Temnothorax* ants in the form of mixed species colonies from a population in Washington State. Mixed colonies consist of workers from two species but only a single species of queen. The following four species were found in mixed colonies: *Temnothorax nevadensis* (Wheeler, 1903), *Temnothorax nitens* (Emery, 1895), *Temnothorax rudis* (Wheeler, 1917), and *Temnothorax rugatulus* (Emery, 1895). Interestingly, all four species mostly occurred free-livingly by themselves and were not found in mixed colonies in the remainder of their distribution range at lower elevation and latitude.

Despite the wide geographic distribution range of these four western *Temnothorax* species, mixed colonies have only been observed in the Pacific Northwest. To characterize the species interactions in mixed colonies, we conducted a series of behavioral experiments and found that populations with mixed colonies show increased aggression towards potential parasites compared to populations where mixed colonies do not occur. We discuss our findings in the context of social parasite evolution (i.e., hypotheses about the origin of dulotic behavior) and discuss whether the variation in ecological environment is associated with the occurrence of mixed *Temnothorax* colonies.

## Methods

### Field site and colony collection

Whole, live colonies of *Temnothorax* ants were collected during June and July of 2014 as well as during July of 2015 ranging from southwestern Arizona to northwestern Washington (Table 1). Colonies were located by thoroughly searching 10 x 10 meter plots. All plots were at least 25 km apart though most were significantly further apart. To find *Temnothorax* colonies, we broke open twigs, pulled apart rotten logs, turned over rocks, and opened rock crevices. Colonies typically ranged in size from 30 to 400 ants, with some very large colonies reaching 1300 individuals (Bengston and Dornhaus 2013). The relatively small colony sizes and the ants’ nesting habit choices allowed for collection of entire colonies, including the queens and brood (Figure A in Supplementary Material). Colonies were collected with an aspirator and stored in separate vials until they were established in artificial nests in the lab.

**Table 1:** Summary of collection site locations, years sampled and if mixed colonies were found or not.

| **Site** | **Location** | **Years Sampled** | **Mixed colonies** |
| --- | --- | --- | --- |
| **1** | **Bear Canyon - White Pass, WA**<br>46.71328, -120.9010 | 2014, 2015 | Present |
| **2** | **Wildcat Creek- White Pass, WA**<br>46.68170, -121.1336 | 2015 | Present |
| **3** | **Palisades - White Pass, WA**<br>46.69483, -121.5465 | 2015 | Present |
| **4** | **Hells Backbone - Boulder, UT**<br>37.96104, -111.5538 | 2014, 2015 | Absent |
| **5** | **Mt. Lemmon - Santa Catalina Mts., AZ**<br>32.39513, -110.6876 | 2014, 2015 | Absent |
| **6** | **Turkey Creek – Chiricauha Mts., AZ**<br>31.91239, -109.2554 | 2014, 2015 | Absent |

Mixed colonies, consisting of workers from at least two species, were first discovered in the colonies collected in 2014 at Bear Canyon, WA (Site 1 in Table 1). To look for additional mixed colonies, ants in all of the collected colonies were visually identified to species in the lab by SEB using Snelling et al. (Snelling et al. 2014).

Censuses of each colony were taken to measure the number of workers, brood, and queens.

To confirm that these mixed colonies were a real phenomenon and not an error in collection (such as accidentally collecting two colonies that nested in close proximity to one another), we repeated our field studies in 2015. At that time, we focused on collecting colonies especially conservatively, which meant only ants from a single cavity or nest chamber were collected and kept alive in separate lab nests. Even with this most conservative collection method, mixed colonies were still found in the sampled colonies. To test whether mixed species nests could be located in adjacent habitats, we added two more collection sites (Wildcat Creek and Palisades, WA; sites 2 & 3, Table 1) in 2015, which were located approximately 10 km from the initial site where mixed colonies were located.

### Laboratory maintenance of live ant colonies

Colonies collected in 2014 were maintained at the University of Arizona in Tucson and colonies collected in 2015 were maintained at the University of Rochester in Rochester, NY. Each colony was established inside a standard artificial nest, which was constructed of a cardboard nest chamber between two glass slides (nest chamber size: 2.75 x 2 cm; Dornhaus et al. 2008, Bengston and Dornhaus 2014). These nests were kept inside a fluon-coated container (10 x 10 cm). Weekly, each colony was fed a small portion of a modified Bhatkar diet: 1 part chicken egg yolk to 1 part raw honey, and multivitamin powder mixed in 5 parts warm water with enough agar powder to form a soft solid when cool (Bhatkar et al. 1970). There was no instance where colonies consumed this entire portion, so colonies likely had *ad libitum* access to food while kept alive in the laboratory. Colonies also had *ad libitum* access to water. Voucher specimens were collected from each colony and were deposited at the Museum of Comparative Zoology at Harvard University (MCZ) and in the Rabeling Lab collection. In addition, vouchers are available upon request from SEB.

### Behavioral analysis

To measure colony-level aggressive behavior, we used a well-established intruder introduction assay (Bengston and Dornhaus 2014, 2015). The ‘intruder’ was a live worker selected from colonies collected in Red Butte Canyon, Utah, which is a site not represented in this study to avoid accidentally using a worker that was related to the focal colony (e.g. a worker from a sister or daughter colony). The intruder was marked with fluorescent powder (Ready Set Glow, product Extreme G12) to increase visibility to the observer and was placed approximately 1 cm into the nest of the focal colony. The introduction event and the following 5 minutes were filmed to facilitate behavioral scoring. To score a colony’s response to the intruder, the interior space of the nest was divided into four quadrants to minimize bias towards behaviours more or less proximal to the intruder and five individuals from each quadrant were observed for 5 seconds every 30 seconds. To further decrease bias from proximity to the intruder, as well as individual worker variation, the average behavioral score was used to represent a colony wide response. All 20 individuals were chosen randomly by the observer at each 30 s interval. This resulted in 10 colony-wide behavioural scores per colony per introduction. The behaviours were scored by the categories listed in Table A of the Supplementary Material, ranging from complete inactivity (score 0) to highly aggressive behaviour (score 6).

Introduction assays were conducted in pairwise combinations of all species represented in the mixed colonies, i.e., including *T. nevadensis*, *T. nitens*, *T. rudis*, and *T. rugatulus*. To detect species-level differences in aggression, all assays were conducted in non-mixed colonies, i.e. those with only one species of workers as determined by visual inspection. Six colonies of each species were selected from each site. The same focal colonies were used for all introductions within a site, with a minimum of 48 hours between introductions. The order in which species were introduced was randomized across colonies to avoid order effects (Bell 2013).

### Statistical analysis

For mixed colonies, we used a Fisher’s Exact Test to look for different frequencies of which species were queenright and which were worker only (Alloway 1980). To evaluate whether aggressive behavior was associated with collection site, focal species, and whether the introduced species was inter- or intraspecific, we used a linear model with aggression as the response variable and with site, colony, focal species, and intruder species as the predictor variables. We performed a stepwise model selection test (both forwards and backwards) to calculate the model with the lowest Akaike’s information criterion (AIC). This allowed us to remove predictor variables that were the least informative to avoid multicollinearity (Derksen and Keselman 1992). We then compared the interaction of these significant predictor variables in two separate generalized linear models: (i) sites with mixed colonies and (ii) sites without mixed colonies. Finally, we used a Tukey’s HSD test to test for variation in aggression between different focal colony species and introduced species. All of the analyses were done using Minitab 17 and R 3.2.2 with RStudio version 0.99.473.

## Results

### Mixed colonies

To evaluate the interspecific interactions in mixed colonies of *Temnothorax* species in the Western United States, we collected a total of 184 colonies (102 colonies in 2014, and 82 colonies in 2015). Across collection sites, 12% of the colonies (n = 22) were identified as mixed-species. Mixed colonies were found exclusively in Washington State (sites 1, 2, and 3; see Table 1) and not at any other site. Among the colonies collected in Washington (total n = 92), 24% were mixed. A total of four species were involved in these mixed-species colonies: *T. rugatulus*, *T. nitens*, *T. nevadensis,* and *T. rudis*. Species pairings with *T. rugatulus* and *T. nitens* were encountered most frequently (77%; Table 2a). However, there was no significant difference between species in their probability of being queenright or consisting only of workers (Fisher’s Exact Probability Test, P = 0.6370). In mixed colonies, there were between 1 and 13 foreign workers, which made up .4% - 17.3% of the total workers in the nest.

**Table 2:** A summary of the mixed colony demographics.

| **Queenright Species** | **Foreign worker species** | **Queen #** | **Colony Size** | **Foreign worker #** |
| --- | --- | --- | --- | --- |
| *T. rugatulus* | *T. nitens* | 4 | 216 | 6 |
| *T. rugatulus* | *T. nitens* | 4 | 61 | 2 |
| *T. rugatulus* | *T. nitens* | 3 | 339 | 4 |
| *T. rugatulus* | *T. nitens* | 3 | 123 | 7 |
| *T. rugatulus* | *T. nitens* | 3 | 198 | 6 |
| *T. rugatulus* | *T. nitens* | 3 | 312 | 8 |
| *T. rugatulus* | *T. nitens* | 3 | 240 | 10 |
| *T. rugatulus* | *T. nitens* | 2 | 282 | 1 |
| *T. rugatulus* | *T. nitens* | 2 | 154 | 3 |
| *T. rugatulus* | *T. nevadensis* | 2 | 244 | 6 |
| *T. rugatulus* | *T. nevadensis* | 1 | 110 | 6 |
| *T. rugatulus* | *T. rudis* | 1 | 110 | 2 |
| *T. nitens* | *T. rugatulus* | 1 | 213 | 7 |
| *T. nitens* | *T. rugatulus* | 2 | 202 | 12 |
| *T. nitens* | *T. rugatulus* | 2 | 194 | 2 |
| *T. nitens* | *T. rugatulus* | 1 | 171 | 1 |
| *T. nitens* | *T. rugatulus* | 1 | 121 | 2 |
| *T. nitens* | *T. rugatulus* | 2 | 111 | 5 |
| *T. nitens* | *T. rugatulus* | 1 | 185 | 6 |
| *T. nitens* | *T. rugatulus* | 1 | 91 | 3 |
| *T. nitens* | *T. rudis* | 1 | 93 | 7 |
| *T. nevadensis* | *T. rudis* | 1 | 75 | 13 |

### Aggressive behavior

The stepwise model selection removed colony as a predictive variable, leaving site, focal species and if the introducer species was inter- or intraspecific in the model (see Table B of the Supplemental Material) . At sites with mixed colonies, the focal species and whether the intruder was inter-specific and intra-specific significant predictor of aggression (two- way ANOVA, focal species P < 0.0001, intruder species P < 0.0001, df = 2876). The most aggressive interactions occurred between *T. rugatlus* and *T. nitens* when either species was the focal species. The next most aggressive interactions were in intraspecific introductions. There was no significant difference in aggression between other species pairings during the other interspecific encounters (Tukey’s HSD test, Figure 1; Table C of the Supplemental Material).

**Figure 1:**
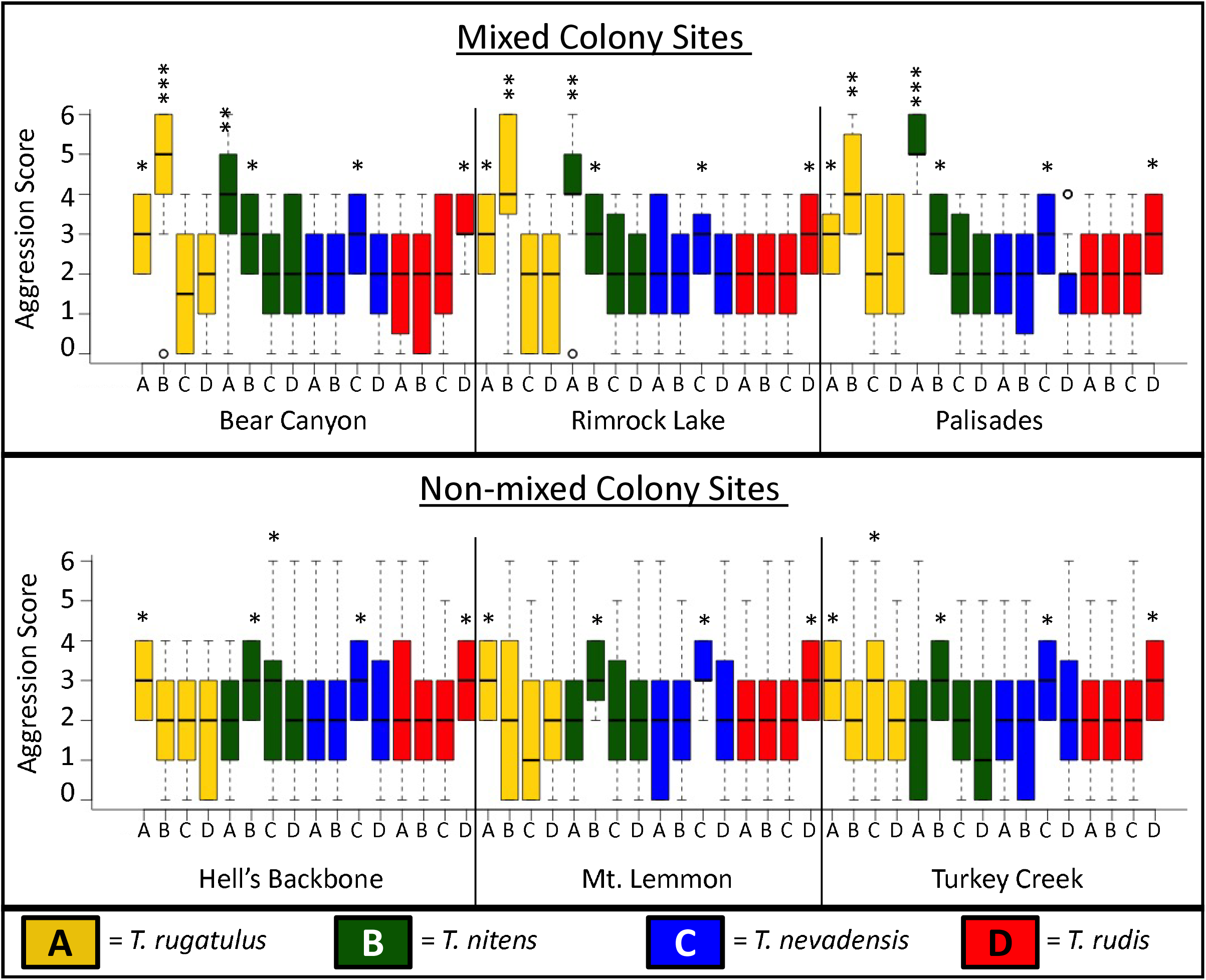
Aggression response to intruder. In colonies from mixed colony sites, the most aggressive interactions were interspecific and occurred between *T. rugatulus* and *T. nitens*. In colonies from non-mixed colony sites, the most aggressive interactions were intraspecific. The aggression score is on the Y-axis and the color of the box indicates the species of the focal colony (e.g. a yellow box represented *T. rugatulus* focal colonies). The letter on the X-axis indicates the species of the intruder. For example, a B under a yellow box indicates a *T. nitens* was introduced into a *T. rugatulus* colony. Stars represent significantly different bins as defined by the Tukey’s HSD post hoc test.

When analyzing colonies from sites consisting of a single species, the only significant predictor of aggression was whether the introduced worker was inter- or intraspecific (two-way ANOVA P < 0.0001, df = 2785 colonies). The most aggressive interactions occurred during intraspecific introductions across all species (Tukey’s HSD test, Figure 1).

## Discussion

Here, we describe the first known instance of mixed-species colonies in four western *Temnothorax* ant species. All colonies only contained a single species of queen, but mixed colonies contained workers of two different species that appeared to be working and living together as in single-species colonies. We investigated this phenomenon across six populations along a 2500 km North-South latitudinal gradient from Arizona to Washington, and mixed colonies were only found in the three most northern sites in Washington State. Four species were involved in this suite of mixed worker colonies, with the most prominent pairing being *T. rugatulus* and *T. nitens*. These two species were equally likely to be queenright.

The presence of mixed colonies with two species of worker and a single species of queen within a single colony is strongly indicative of social parasitism. So far, observations of mixed colonies, where the brood of two species is mingled and communally cared for (*sensu* Wasmann 1891), is characteristic of socially parasitic interactions. In contrast, species interactions in compound nests, where two species share a nest space but keep their brood separately, may range from mutualuistic to parasitic/predatory inteactions (Hölldobler and Wilson 1990b; Kaufmann et al. 2003; Gray et al. 2018). Because all four species that can be part of a mixed colony are also encountered free-living, it appears that the observed interactions constitute a form of facultative social parasitism. Additionally, all of these species can found new colonies independently (Rüppell et al. 1998), SEB, unpublished data), meaning they are not dependent on another species or colony to begin worker or alate production. This is an important distinction between obligate and facultative social parasitism, as obligate social parasite queens cannot found new colonies independently.

However, the simple presence of a mixed colony does not clearly define the life history strategy of the social parasite. Our observations are consistent with (a) dulosis, where workers of the queenright species raid neighboring colonies to steal their brood, or (b) temporary social parasitism, where the parasitic queen invades a host colony, kills the resident queen, and usurps the worker force to raise her offspring. Inquilinism is not a likely explanation for the pattern we find here in the Western *Temnothorax* species because: (i) many inquiline social parasites are host queen tolerant, and therefore, queens of both host and parasite would be expected inside the same colony; however, we only ever found the queen of a single species in mixed colonies. (ii) Most inquiline species do not produce a worker caste, and in that case, only the workers of the host species would be detected inside the colony. In contrast, we observed the workers of two species in mixed colonies.

We suggest that our observations likely represent a case of facultative dulosis rather than temporary social parasitism. Our argument is based on the following three lines of evidence. First, queens of temporary social parasites specialize in entering established host colonies, killing the resident queen, and utilizing her workforce to raise their own brood (Heinze and Keller 2000). However, in all mixed colonies, the workers belonging to the same species as the queen were vastly the majority over the non- queenright species, which is more consistent with dulotic brood stealing behavior (Table 2b). If a colony was in the process of a slow turnover of species, as in temporary social parasitism, it would be surprising that by random chance we collected colonies all in the late stage of this process, rather than finding a mix of host to parasite ratios. Additionally, the queenright species was often polygynous, meaning there were multiple queens in the colony (Table 2b).

Second, the behavioral results are consistent with facultative dulosis. Aggressive territoriality behavior has been repeatedly highlighted as a prerequisite behvavior for dulosis to evolve as well as a defense against parasitism (Topoff 1990; Huang and Dornhaus 2008). In this study, we show that *T. rugatulus* and *T. nitens* are significantly more aggressive towards each other in mixed colony sites compared to sites where mixed colonies have not been found; perhaps signaling that this increased aggression is a component of an escalating arms race against facultative dulosis, which is consistent with observations on eastern *Temnothorax* colonies that have been threatened by dulotic species (Pamminger et al. 2012). In comparison, colonies from both mixed colony and non-mixed colony sites show intermediate aggression towards conspecifics. *Temnothorax rugatulus* and *T. nitens* from mixed colony sites do not show increased aggression towards the other *Temnothorax* species tested here (*T. rudis* or *T. nevadensis*).

Third, the ecology at the Washington sites, where mixed colonies have been found, is conducive of territorial contests, which could lead to facultative dulosis in *Temnothorax* ants. Previous studies have shown that *Temnothorax* at the northern sites are under higher competition for nest sites, meaning that more of the potential nest sites are occupied than at southern sites (Bengston and Dornhaus 2015). Because *Temnothorax* are obligate cavity dwellers, this likely limits their population size. Additionally, due to differences in where potential nest sites are located, colonies in the northern sites are more closely spatially clustered (Bengston and Dornhaus 2015), perhaps suggesting colonies are more likely to encounter one another and increasing the opportunity for territorial contests.

Territorial contests were repeatedly hypothesized as a first step in the evolution of dulotic behavior where workers have maintained the ability to perform normal colony tasks without the need for host workers (Sakagami and Hayashida 1962; Wilson 1971; Mori and Moli 1988; Mori et al. 2001). Alloway (Alloway 1980) observed in the lab that territorial contests resulted in mixed colonies of the Eastern US species *Temnothorax ambiguus* (Emery, 1895) and *Temnothorax longispinosus* (Roger, 1863). Both species are hosts of social parasites but neither of these species is an obligate social parasite, and brood stealing seems to be a facultative, opportunistic phenomenon (Alloway 1980). In fact, evolutionary transitions to dulotic behavior occurred at least three times independently in *Temnothorax* ants (Prebus 2017). In North America, the dulotic species *Temnothorax americanus* (Emery, 1895), *Temnothorax duloticus* (Wesson, 1937), and *Temnothorax pilagens* Seifert *et al*., 2014 parasitize *Temnothorax ambiguus*, *Temnothorax curvispinosus* (Mayr, 1866), and *Temnothorax longispinosus* (Wilson 1975; Alloway 1979; Seifert et al. 2014). Alloway (1980) additionally suggested that *T. ambiguus* and *T. longispinosus* are “highly preadapted for social parasitism” and “all that would seem required to transform one of these infrequent facultative brood stealers [updated segments] into a frequent social parasite would be a few quantitative changes in the frequencies with which workers engage in certain behaviors” (1980, p 255). These certain behaviors may simply be initiating territorial disputes and/or raiding behavior after a successful territorial defense. The observed increased aggressive behavior may be particularly beneficial during territorial contests. In conclusion, the combined direct and indirect evidence suggests that the mixed colonies of western *Temnothorax* species constitute a case of facultative dulotic social parasitism.

Finally, this study brings attention to the possibility of undetected “intraspecific parasitism” (Hölldobler 1976; Foitzik and Heinze 1998). This phenomenon is similar, functionally, to interspecific parasitism. For example, harvester ant colonies (*Pogonomyrmex rugosus*) have been observed raiding smaller colonies for brood.

Microsatellite data showed that 43% of colonies had foreign workers, with the proportion of foreign workers ranging from 6-28% of the foragers sampled (Gadau et al. 2003).

Territorial aggression within species, then between competing species (all studied *Temnothorax* species compete for the same, limiting, nest sites) may provide one evolutionary pathway towards obligate, interspecific dulotic social parasitism in myrmicine ants (Alloway 1980; Pollock and Rissing 1989; Huang and Dornhaus 2008). Interestingly, dulosis evolved convergently and under different ecological circumstances in the distantly related subfamily Formicinae and in the *Formica sanguinea* group, predatory behavior gave rise to facultative dulotic bahvior (Borowiec et al. 2021).

Understanding how intra- and inter-specific social parasitism can change within-colony relatedness is an important next step, as it has significant implications for the “super organism” concept and our assumptions about worker origin, even if a colony appears to be monogynous.

