## Supplementary material for "The discovery of mixed colonies in *Temnothorax* ants supports the territoriality hypothesis of dulotic social parasite evolution in myrmicine ants"

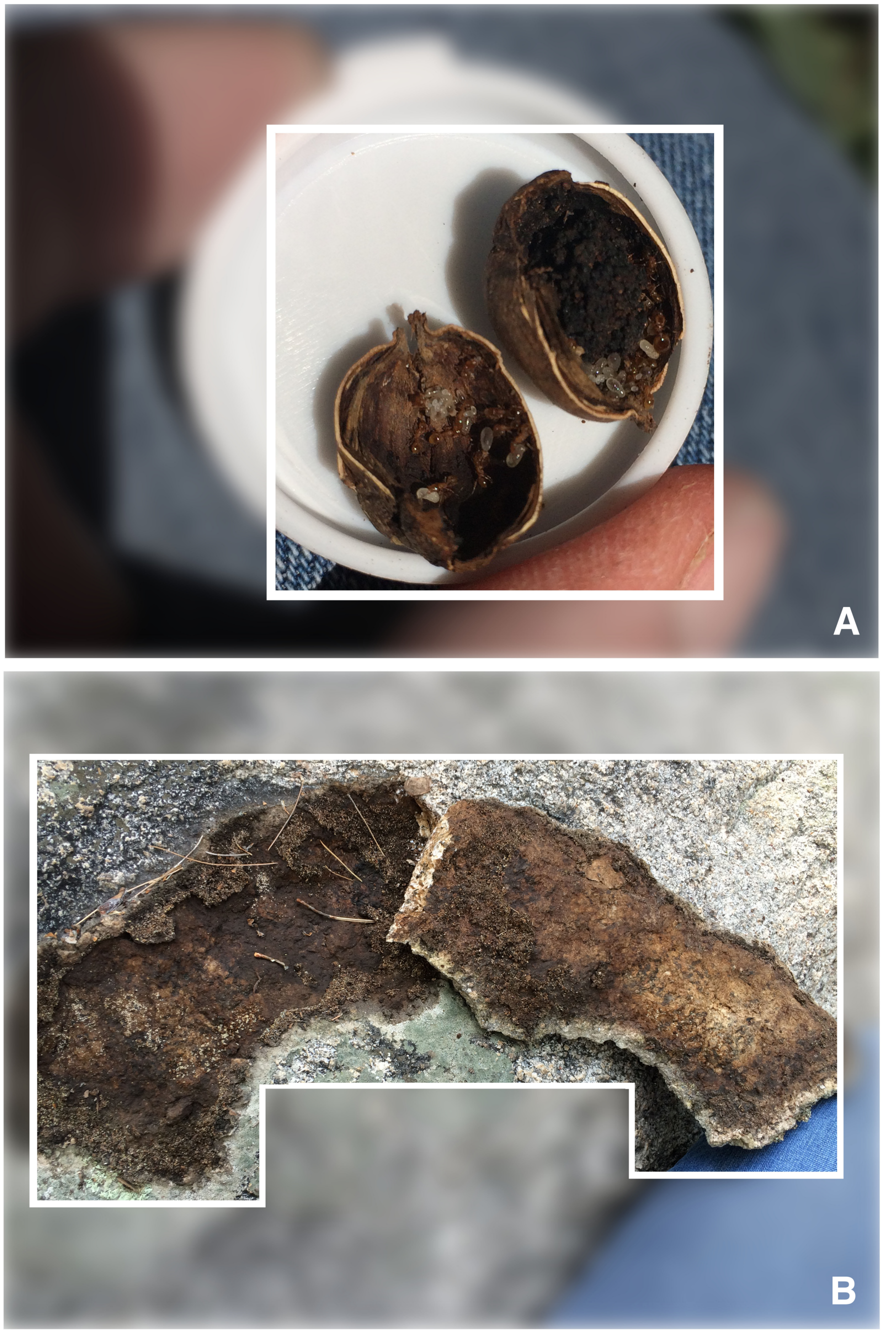

Figure A: Example nest sites. Frame A shows a colony of *Temnothorax nitens* inside an acorn. Frame B shows a *Temnothorax rugatulus* nest between two rocks.

| Score | Example Behaviors |
|---|---|
| 0 | Inactive, self grooming, antennating nest mates |
| 1 | Grooming intruder, crouching |
| 2 | Exit nest, move brood away from intruder |
| 3 | Antennal drumming, climbing on intruder |
| 4 | Extending legs of intruder, mandibular fencing |
| 5 | Carrying intruder towards nets entrance, dragging intruder |
| 6 | Stinging or biting intruder |

Table A: Behaviors and their associated scores used to quantify behavioral responses to an intruder.

| Model Residuals: | | | | |
| --- | --- | --- | --- | --- |
| **Min** | **1Q** | **Median** | **3Q** | **Max** |
| -2.8231 | -1.2456 | 0.0907 | 1.289 | 3.9786 |
| **Coefficients** | | | | |
| | **estimate** | **std. Error** | **t-value** | **p-value** |
| **Intercept** | 3.2320 | 0.0759 | 42.5810 | < 0.0001 |
| **site** | -0.0675 | 0.0114 | -5.9210 | < 0.0001 |
| **focal species** | -0.1121 | 0.0174 | -6.4370 | < 0.0001 |
| **intro. species** | -0.1174 | 0.0174 | -6.7400 | < 0.0001 |
| **Residual Standard Error** | | | | |
| 1.478, df = 5756 | | | | |
| **Adjusted R2** | | | | |
| 0.0202 | | | | |

**Table B:** The model residuals of the stepwise selected model, which determined the best predictors for aggression score were collection site, the focal species being observed, and the species of the intruder.

| B1 | | | | | |
|---|---|---|---|---|---|
| Sites with mixed colonies | | | | | |
| **Focal Species** | **Introduced species** | | | | **Mean** |
| *T. nitens* | *T. rugatulus* | A | | | 4.48 |
| *T. rugatulus* | *T. nitens* | A | | | 4.46 |
| *T. nevadensis* | *T. nevadensis* | | B | | 3.12 |
| *T. rudis* | *T. rudis* | | B | | 2.97 |
| *T. nitens* | *T. nitens* | | B | | 2.94 |
| *T. rugatulus* | *T. rugatulus* | | B | | 2.94 |
| *T. nitens* | *T. rudis* | | | C | 2.17 |
| *T. nitens* | *T. nevadensis* | | | C | 2.11 |
| *T. rugatulus* | *T. nevadensis* | | | C | 2.08 |
| *T. rudis* | *T. rugatulus* | | | C | 1.99 |
| *T. rugatulus* | *T. rudis* | | | C | 1.98 |
| *T. nevadensis* | *T. rudis* | | | C | 1.97 |
| *T. nevadensis* | *T. rugatulus* | | | C | 1.96 |
| *T. rudis* | *T. nevadensis* | | | C | 1.92 |
| *T. nevadensis* | *T. nitens* | | | C | 1.90 |
| **B2** | | | | | |
| Sites without mixed colonies | | | | | |
| **Focal Species** | **Introduced species** | | | | **Mean** |
| *T. rudis* | *T. rudis* | A | | | 3.16 |
| *T. rugatulus* | *T. rugatulus* | A | | | 3.07 |
| *T. nitens* | *T. nitens* | A | | | 3.03 |
| *T. nevadensis* | *T. nevadensis* | A | | | 2.98 |
| *T. rudis* | *T. nevadensis* | | B | | 2.20 |
| *T. nitens* | *T. rudis* | | B | | 2.19 |
| *T. nevadensis* | *T. rugatulus* | | B | | 2.16 |
| *T. nevadensis* | *T. rudis* | | B | | 2.08 |
| *T. nevadensis* | *T. nitens* | | B | | 2.07 |
| *T. nitens* | *T. rugatulus* | | B | | 2.02 |
| *T. rugatulus* | *T. nitens* | | B | | 1.99 |
| *T. rudis* | *T. rugatulus* | | B | | 1.94 |
| *T. rugatulus* | *T. nevadensis* | | B | | 1.92 |
| *T. nitens* | *T. nevadensis* | | B | | 1.88 |
| *T. rugatulus* | *T. rudis* | | B | | 1.88 |

Table C: The connecting letters report of the Tukey's HSD test comparing the aggression associated with different pairwise combinations of focal and intruder species, as seen in

Fig. 1. B1 shows the values associated with the mixed colony sites; B2 shows the sites without mixed colonies.
